# Deciphering the role of brainstem vestibular-related inhibitory networks in shaping postural reflexes in the Xenopus tadpole

**DOI:** 10.1101/2025.11.16.688675

**Authors:** Léandre Lavenu, Mathilde Pain, Gabriel Barrios, Laura Cardoit, Marie Boulain, Alexia Duveau, Hervé Tostivint, François M. Lambert, Pascal Fossat, Didier Le Ray

**Affiliations:** Univ. Bordeaux, CNRS, INCIA, UMR 5287, F-33000 Bordeaux, France; Univ. Bordeaux, CNRS, IMN, UMR 5293, F-33000 Bordeaux, France; MNHN, CNRS, PhyMA, UMR7221, F-75000 Paris, France

**Keywords:** Vestibular networks, Gaba, Glycine, Posture, Xenopus

## Abstract

Brainstem vestibulospinal (VS) nuclei generate excitatory commands in response to multi-modality sensory integration, to activate specific spinal networks in order to generate adapted postural reflexes. Comparably organized in bilateral nuclei with both ipsi- and contralateral pathways in all species, excitatory VS projections alone fail to explain the mostly unilateral reflex responses typically observed. In the *Xenopus laevis* tadpole, we describe secondary vestibular neurons of inhibitory nature, and the synaptic contacts they make on VS neurons. Then, using a brainstem/spinal cord in vitro preparation we show that the spinal responses evoked by galvanic vestibular stimulation are shaped by both commissural and local inhibitory brainstem networks. We further show that a complex interaction between GABAergic and glycinergic inhibitory networks regulate VS neuron excitability and, consequently, the expression of the spinal response. Our data reveal that while excitatory VS neurons execute the neural score, inhibitory neurons in the central vestibular system coordinate and modulate the overall performance.

## Introduction

The control of posture is a complex sensory-motor mechanism that is a common to all animals, whatever their ecophysiology (Cattaert & Le Ray, 2001; MacKinnon, 2018; Le Ray & Guayasamin, 2022). In vertebrates, the vestibulospinal (VS) system plays a pivotal role, by integrating various sensory afferent and motor efferent modalities to generate adapted postural commands (Cullen, 2019). Comparing the VS organization across species proved considerable conservation among vertebrates (Glover, 2001; Díaz & Glover, 2002; Straka & Baker, 2013).

Vestibular sensory neurons encode head accelerations that are perceived both by semicircular canal (head rotations) and otolithic (linear acceleration) organs, and project onto secondary vestibular neurons hosted in central vestibular nuclei (CVN) bilaterally distributed in the brainstem. Two of these bilateral nuclei are more specifically involved in the control of neck and body posture in space. The lateral vestibulospinal tract nucleus (LVST) contains VS neurons that project exclusively into the ipsilateral spinal cord. Besides, a more medial nucleus contains neurons that project either only contralaterally in lower vertebrates (Tangential nucleus, or TAN) or bilaterally in higher vertebrates (medial vestibulospinal tract nucleus; for a review: Glover, 2001). Spinal targets of VS neurons also seem largely conserved in vertebrates and consist both of motoneurons and various segmental interneurons (Shinoda *et al*., 2006; Kasumacic *et al*., 2010; Lambert *et al*., 2016; Olechowski-Bessaguet *et al*., 2020; Zhu *et al*., 2020; Barrios *et al*., 2024). Thus, as considered to date, the VS sensory-motor system consists of a cascade of excitatory commands, from peripheral vestibular sensory neurons to central VS neurons, down to spinal neurons and postural muscles. However, the existence of both ipsilaterally and contralaterally projecting VS neurons that are likely activated simultaneously by unilateral vestibular inputs should result in bilateral spinal responses, incompatible with an adapted postural correction. In this context, although hitherto barely investigated, intrinsic inhibitory networks likely play major roles in shaping accurate vestibulospinal reflexes.

Local and commissural vestibular inhibitory neurons have been identified long ago within CVN, and their impact on vestibulo-ocular reflexes (VOR) was largely documented (Spencer & Baker, 1992; Reichenberger *et al*., 1997; Straka & Dieringer, 2004; Gliddon *et al*., 2005; Malinvaud *et al*., 2010). In addition, numerous studies in frogs demonstrated that secondary vestibular neurons, although not all identified functionally, received massive vestibular sensory-related inhibitory inputs (Straka & Dieringer, 1996, 2000; Straka *et al*., 1997; Biesdorf *et al*., 2008). In contrast, very little is known about their putative contribution to VS reflexes, and the most relevant data come from studies on the antigravity system. Antigravity postural reflexes originate from vestibular otolithic signals, the integration of which is modulated by both local and commissural vestibular inhibitions (Uchino, 2004; Gliddon *et al*., 2005), and manipulating pharmacologically these inhibitory systems was shown to alter body support (e.g., Pompeiano *et al*., 1993). Still, it remains unknown whether vestibular commissural and local inhibitory networks participate in other postural functions (such as body orientation, balance…) and, thus, globally controls how vestibular postural commands are organized.

This fundamental question was addressed in the *Xenopus* tadpole, in which we have recently shown that horizontal semicircular canal signals were the most efficient to generate reliable spinal reflexes, and have detailed the underlying anatomo-functional excitatory pathways. To summarize, vestibular inputs excite ipsilateral VS neurons from both the LVST and TAN nuclei, which in turn mono- and oligosynaptically activate motoneurons in the ipsilateral and contralateral spinal cord, respectively (Barrios *et al*., 2024). Here, we describe secondary vestibular inhibitory neurons in the vestibulospinal nuclei, and report the existence of numerous inhibitory synapses on VS neurons. On a reduced brainstem/spinal cord preparation with intact vestibular endorgans we combined electrophysiology and pharmacology to demonstrate that brainstem inhibitory systems shape postural reflexes recorded from spinal ventral roots. We reveal a complex interaction between glycinergic and GABAergic secondary vestibular neurons, and their direct and indirect impact on VS neurons. We further show that integrity of vestibular commissural inhibition is mandatory for functional vestibular-evoked spinal reflexes to be produced. Our data clearly highlight the prominent role of brainstem inhibitory networks in shaping accurate vestibular-evoked spinal postural responses.

## Materials and methods

### Animals

*Xenopus laevis* embryos were obtained from adult frogs hosted the Institute of Neurodegenerative Diseases (University of Bordeaux; certification #B33-063-942). Then, embryos were bred until stages of interest, in 12/12h dark/light cycle at 22°C, in INCIA facility (certification #B33-063-923). Developmental stages were determined based on external characteristics (Niewkoop and Faber, 1956). All experiments were conducted on premetamorphic tadpoles at stage 52-55, according to European ethic policy and after agreement from local and national ethic committees (APAFIS #38744-2022092911238633 v10).

### Vestibulospinal neuron retrograde labelling

For neuroanatomical characterization, retrograde tracers were applied as previously extensively described in *Xenopus* tadpoles (Barrios *et al*., 2024). Briefly, after forebrain ablation and dorsal skin and cartilage removal under deep anesthesia in buffered 0.05% tricaine methanesulfonate (MS-222) brainstem and spinal cord were isolated *in vitro*. Then, a small incision was made unilaterally in rostral spinal segments 2-3, tracer crystal was directly applied at the ventral part of the section, and preparations were kept in oxygenated (carbogen: 95% O2/5% CO2) saline (120 mM NaCl, 2.5 mM KCl, 30 mM NaHCO_3_, 11 mM glucose, 5 mM CaCl_2_, 1 mM MgCl_2_, pH 7.4) for minimum 5h to allow retrograde dye migration up to the vestibulospinal cell bodies in the brainstem.

Either Alexa Fluor Dextran (AFD) 647 or Neurobiotin (NB) were used depending on the subsequent immunohistochemical treatment: AFD were used when classical immunofluorescence treatment was subsequently applied, while NB and later streptavidin 647 revelation was used in case of in situ hybridization.

After migration, preparations were pinned down in a Sylgard-lined Petri dish and fixed overnight at 4°C in 4% paraformaldehyde (PFA). The day after, tissues were rinsed twice 15 minutes in phosphate buffer saline (PBS: 137mM NaCl, 2.7 mM KCl, 10mM Na_2_HPO_4,_ 1.8mM KH_2_PO_4_), embedded in Tissue-Tek (VWR-Chemicals), frozen (−45°C) in isopentane, and finally stored at −80°C until later treatment.

For patch-clamp experiments (see below) tetramethyl-rhodamine-coupled dextran crystals 3kD (RDA, Life Technologies) were similarly used (although without post-fixation) to retrogradely label VS neurons priorly to the recordings.

### *In situ* hybridization (ISH)

Complementary DNA fragments encoding for *X. laevis* GAD65 (XM_041565850.1) and SLC6A5 (XM_041589711.1) were amplified by RT-PCR (for primer sequence, see Table 1) from adult *X. laevis* brain and spinal cord RACE-ready cDNA as previously described (Bougerol *et al*., 2015), subcloned into pGEM T-Easy vector (Promega) and sequenced for confirmation. Purified cDNA fragments (585 bp for GAD65, and 886 bp for SLC65) were used to synthesize digoxigenin-labelled sense and anti-sense cRNA probes *in vitro* with RNA labelling Mix (Sigma Aldrich) following the manufacturer’s instructions.

**Table 1:**
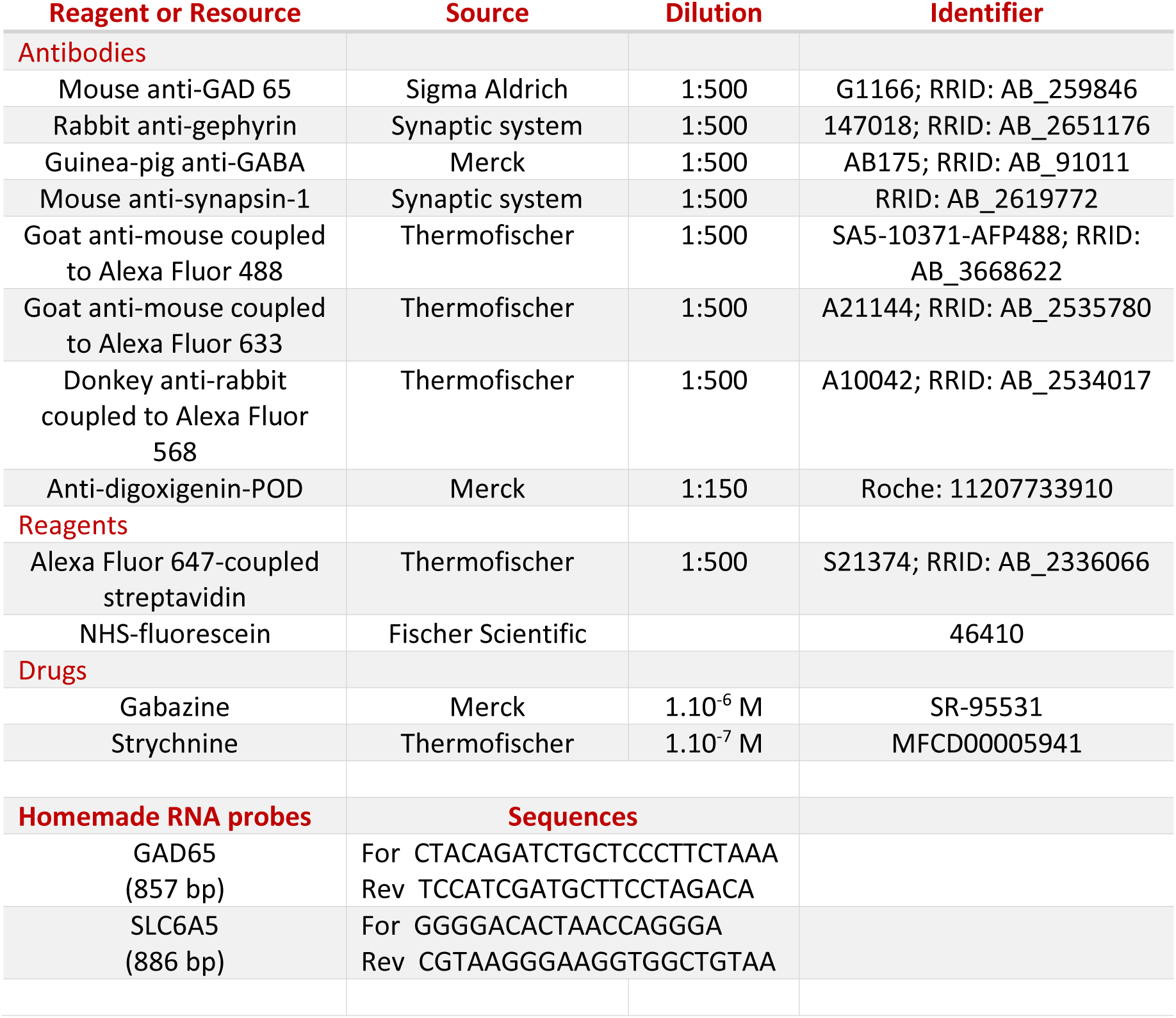
Antibodies and reagents.

Before starting the ISH protocol, three solutions were: a 20x saline sodium citrate buffer (SSC) containing 175.3g NaCl, 88.2g Na3C6H5O7 for 1L of milliQ water, pH 7 + 1mL DEPC incubated overnight under agitation at room temperature, a maleate tampon solution (100mM maleic acid, 250mM NaCl, pH=7.5), and a blocking reagent solution (5g of blocking reagent powder in Maleate tampon solution for 50ml, warmed at 60°C for 1h).

16-µm cryostat slices were made from NB-labeled brainstems, positioned on slides, dried, and then washed four times 15 minutes in PBS-Tween solution at room temperature. Thereafter, in a fume cupboard, slides were incubated in a primary hybridization solution (50% Formamide, 5x SSC, 50mg/mL heparin, 0.5mg/mL yeast tRNA, 0.1% Tween-20, 9.2mM citric acid; pH 6) at 64°C for 1h30 in humid chamber. Next, slides were incubated in homemade digoxigenin-tagged probes (priorly denatured at 80°C in the same primary hybridization solution) at 64°C overnight in humid chamber.

The day after, slides were washed at 65°C in 3 consecutive 20-min baths of respectively 75% secondary ISH solution (50% formamide, 5x SSC, 0.1% Tween-20) +25% 2x SSC, 50% ISH+50% 2x SSC, and 25% ISH+75% 2x SSC. Afterwards, slides were washed 20 minutes in 2x SSC, then twice 30min in 0.05x SSC at 65°C, and rinsed for 5min in 50% 0.05x SCC+50% PBS-Tween solution, and twice 5min in PBS-Tween at room temperature. Thereafter, slides were covered with a blocking reagent solution during 2h at room temperature. The specificity of the hybridization procedure was verified by incubating sections with the sense riboprobes with which only background signals typical of this type of chemical reaction could be observed (not illustrated).

### Immunofluorescence treatment

Immunolabelling: Inhibitory synapses have been targeted using primary antibodies directed either against Gad65 (Sigma Aldrich, 1:500 dilution), a presynaptic GABA terminal protein (to label only GABAergic synapses), or against gephyrin (Synaptic system, 1:500), a postsynaptic inhibitory synapse protein (to label all inhibitory synapses). Briefly, 20-µm slices were cut from AFD 647-labeled brainstems using a cryostat (Leica 3000), then positioned on 52°C gelatin-coated slides (Superfrost+; Epredia) for 5-10 minutes, and stored at −20°C until later immunotreatment. First, slides were immerged in PBS-BSA-Triton solution (1% Bovine Serum Albumin, 0.3% Triton-X500) during two hours to saturate and permeate tissues. After washing the slides three times 15 minutes in PBS, anti-GAD-65 or anti-gephyrin primary antibodies was applied overnight, in dark wet chamber at 4°C. Thereafter, antibodies were washed out in PBS (3×15 min), and the corresponding secondary antibody (see Table 1) then applied for 2h in dark wet chamber at room temperature.

Fluorescent ISH (FISH): Inhibitory neurons hybridized with digoxigenin-tagged probes were labeled using an anti-digoxigenin-POD antibody conjugated with horse radish peroxidase, applied overnight at 4°C in humid chamber, and subsequent revelation with NHS-fluorescein (see Table 1). Consecutively, slides were also incubated in streptavidin 647 in order to make visible the VS neurons retrogradely labeled with NB.

At the end, slides were washed 3 times in PBS, rapidly dipped into water and mounted in Fluoromount solution (Southern Biotech).

### Image acquisitions and analysis

Fluorescence signals were acquired using a Zeiss confocal microscope LSM900 equipped with 488,543,633 nm lasers. Global slides acquisitions were performed with a 10X/0.45 dry objective, while multi-images confocal Z-stack acquisitions were acquired with both a 20X/0.8 dry objective and a 40X/1.4 oil objective with 1µm step intervals. Final images were processed with artificial fluorescent colors using Fiji (https://fiji.sc/). Both signal apposition and co-localization were identified and counted using a homemade script based on the Fiji macro DiAna (Gilles *et al*., 2017). Briefly, neurons were identified by their size and circularity on 20-µm confocal image stacks, under user control in order to avoid false negatives, and the number of contacts on the 3D cell surface was subsequently counted.

### *In vitro* brainstem/spinal preparations for electrophysiology

Brainstem and spinal cord reduced preparations with intact otic capsules were dissected and recorded as previously detailed (Barrios *et al*., 2024). Briefly, animals were anesthetized in buffered MS-222 solution, and the forebrain rostral to the mesencephalon was rapidly removed for ethic purpose. Then, dorsal skin and muscles were removed, together with the dorsal cartilage covering the CNS. Next, spinal ventral roots (Vr) were gently separated from the remaining cartilage and muscles, and the ensemble head/brainstem/spinal cord was pinned down a recording chamber (Sylgard-lined Petri dish) and continuously perfused with oxygenated saline at 18°C. A thin, waterproof Vaseline wall was built around and above the first spinal segment in order to perfuse separately the brainstem from the spinal cord, allowing the application of pharmacological drugs restricted to the brainstem.

Ventral roots were recorded from several spinal segments using glass suction electrodes (TF150; Harvard apparatus), the resulting extracellular electrophysiological signals directed to a Model-1700 amplifier (AM-systems) and digitized at 10kHz through a Micro-1401 interface (Cambridge Electronic Design). All electrophysiological signals were acquired, stored, and analyzed using Spike2 software (CED).

### Galvanic vestibular stimulation (GVS)

In order to mimic the activation of the vestibular endorgans that are the most stimulated during natural swimming in tadpoles (Barrios *et al*., 2024), GVS was delivered using two coated iron electrodes placed in contact with the otic capsule on both sides, vertical to the horizontal canal sensory cells. GVS consisted of 10 cycles of sinusoidal current (0.1-1Hz/max ±1mA) applied simultaneously but in phase opposition through the left and right otic capsules, close to the canal cupula. 2s/±0.5mA stimulation proved optimal to generate reproducible reflex responses in spinal ventral roots, and all data reported here were obtained using such GVS parameters.

### Central vestibulospinal nuclei stimulation

In order to get rid of putative modulation of sensory signals by antagonists, in distinct sets of experiments direct electrical stimulation was delivered to each central vestibular nucleus (CVN) separately, using a unipolar tungsten electrode (0.1 MΩ; Microprobe) connected to a Model 2100 stimulator (AM-systems). Stimulation consisted of 200-µs current pulses, the intensity of which was set, for each single CVN in each preparation, as the minimal intensity (14 to 70µA) able to evoke in control saline spinal responses within the first 20ms in >90% of trials (see Barrios *et al*., 2024). Then, stimulation was delivered 5 times, with a 1-min interval, and both the response occurrence (0-5) and the strength of spinal response were measured in control saline and under inhibitory receptor antagonist perfusion on the brainstem. Response strength was estimated as the area of the integrated burst (signal was first rectified and smoothed) and the mean firing during burst.

### Patch-clamp recordings

Isolated brainstem/spinal cord preparations with RDA-labeled VS neurons (see above) were partially included by the ventral side in a low melting temperature Agar block (4%; Sigma Aldrich). Using cyanoacrylate, the Agar block was then cemented with the brainstem dorsal-side-up in a vibratome cutting chamber (Leica VT 1000S) filled with oxygenated sucrose solution at 4 °C (1.15mM NaH2PO4, 2mM KCL, 7mM MgCl2, 0.5mM CaCl2, 26mM NaHCO3, 11mM glucose, 205mM sucrose). The dorsal part of the brainstem was removed using a vibrating microslicer to expose VS nuclei. The ventral part of the preparation was then held in oxygenated physiological saline at room temperature for 30 minutes until use.

Preparations were transferred to a recording chamber under an upright microscope (Nikon Eclipse FN1) and continuously perfused with oxygenated physiological solution. RDA-positive VS neurons were visually identified by their anatomical position throughout rhombomeres 4 to 6, using a 570-nm fluorescence system (Dual Optoled Supply) and a 40x water-immersion objective. Microelectrodes were pulled from borosilicate glass capillaries GC120F-10 (1.2 OD x 0.69×100 L mm) on a pipette puller (Sutter Instrument Co. Model p.97). Electrophysiological signals were acquired with 5-8 MΩ patch pipettes filled with homemade intrapipette solution (115 mM K-gluconate, 2 mM MgCl2, 2 mM EGTA, 10 mM HEPES, 2 mM MgATP, 0.2 mM NaGTP). Whole-cell configuration patch clamp recordings of 27 RDA-positive VS neurons were conducted under voltage and current clamp conditions with an Axoclamp 2A amplifier (Axon Instruments). Signals were digitized at 10KHz using a A/D converter interface (Digidata 1440; Axon Instruments) and acquired on computer with Pclamp10 software (Molecular Devices, Berkshire, UK). Series resistance (typically 12-20 mΩ) was measured throughout the recordings, and only stable neurons (changes <10%) were kept for later analysis.

### Drugs

The fast GABAergic neurotransmission was blocked using the GABA-A receptor antagonist gabazine (1 µM; Merck; see Table 1), which was previously dissolved in distilled water in 1-ml aliquots stored at −20°C. The glycine receptor antagonist strychnine (100 nM; Thermofischer Scientific; see Table 1) was used to disrupt glycinergic transmission. Strychnine had been previously dissolved in distilled water in 1-ml aliquots stored at −20°C. Unfortunately, even at these low concentrations none of the drugs could be washed out from the brainstem/spinal cord preparation during in vitro experiments, so each drug was applied separately on different sets of animals.

### Statistical analysis

GVS-evoked reflex responses were analyzed with circular statistics using Oriana 2 software (Kovacs). For a given ventral root, all phases of all GVS cycle-related responses were calculated as the delay to reach the response peak referred to the cycle duration, and the mean phase response for the spinal root was calculated as the mean vector angle for the whole stimulation sinusoid. Only significant mean phase values were considered, i.e., when mean vector length was larger than 0.5 and the Rayleigh test (which test for distribution uniformity) was significant. Then, all significant mean phase values for a given root were pooled over several sinusoid stimulations, and circular graphs were generated to illustrate the population mean response and to perform statistical comparisons. Watson-Williams F-test was used to compare mean vectors in control and drug conditions. Significance threshold was set at 0.05.

Intracellular PPSI recordings and CVN electrical stimulation-evoked spinal responses were analyzed using Prism 10 software (Graphpad). PPSI amplitudes, durations and frequencies were compared with ANOVA tests. Results are expressed as mean ± SEM. Changes in minimal stimulation intensity, response occurrence, and changes in response strength were tested with either a Wilcoxon paired t-test (two conditions) or one-way ANOVA (three conditions), with a significance threshold of 0.05.

## Results (1736 mots)

### GABAergic and glycinergic contacts on identified VS neurons

Retrograde AFD labeling of VS neurons (LVST: n=152; TAN: n=176; n=3 animals) from a proximal spinal segment was combined to immune detection of gephyrin, a synaptic protein found in all ionotropic inhibitory synapses (Vitanova *et al*., 2004), and Gad65, specific of GABA-mediated synapses (Fig. 1A-D). Most of VS neurons in LVST (73.0 %) and TAN (73.9 %) displayed a marked gephyrin immunolabeling around soma and dendrites (example of LVST VS neuron in Fig. 1B), demonstrating that they received abundant inhibitory synapses (Fig. 1C). Conversely, a significant proportion of LVST (27.0 %) and TAN VS neurons (26.1 %) showed no inhibitory contacts (Fig. 1C), and likely only received excitatory inputs, as previously reported from unidentified secondary vestibular neurons in a terrestrial frog (Straka *et al*., 1997). Interestingly, 39.5 % of identified VS neurons in LVST and 54.6 % in TAN (Fig. 1C-D) showed both gephyrin and Gad65 immunolabeling, either colocalized or not, suggesting the presence of both GABA-mediated and non-Gaba-mediated (likely glycinergic; see Straka *et al*., 1997) inhibitory synapses. The remaining proportion of gephyrin-positive VS neurons likely received inhibitory inputs of exclusively glycinergic nature (LVST: 33.5 %; TAN: 19.3 %). In VS neurons displaying by both GABAergic and putatively glycinergic contacts (Fig. 1D), a large majority of non Gad65-associated gephyrin puncta were found in the two nuclei (LVST: 6±2 Gad65 and 15±3 non-Gad65; TAN: 12±2 Gad65 and 31±4 non-Gad65; Wilcoxon tests, p<0.001). Taken together, these results suggested that direct putatively glycinergic inhibition might be dominant on VS neurons in *Xenopus* tadpoles, especially on TAN VS neurons.

**Figure 1:**
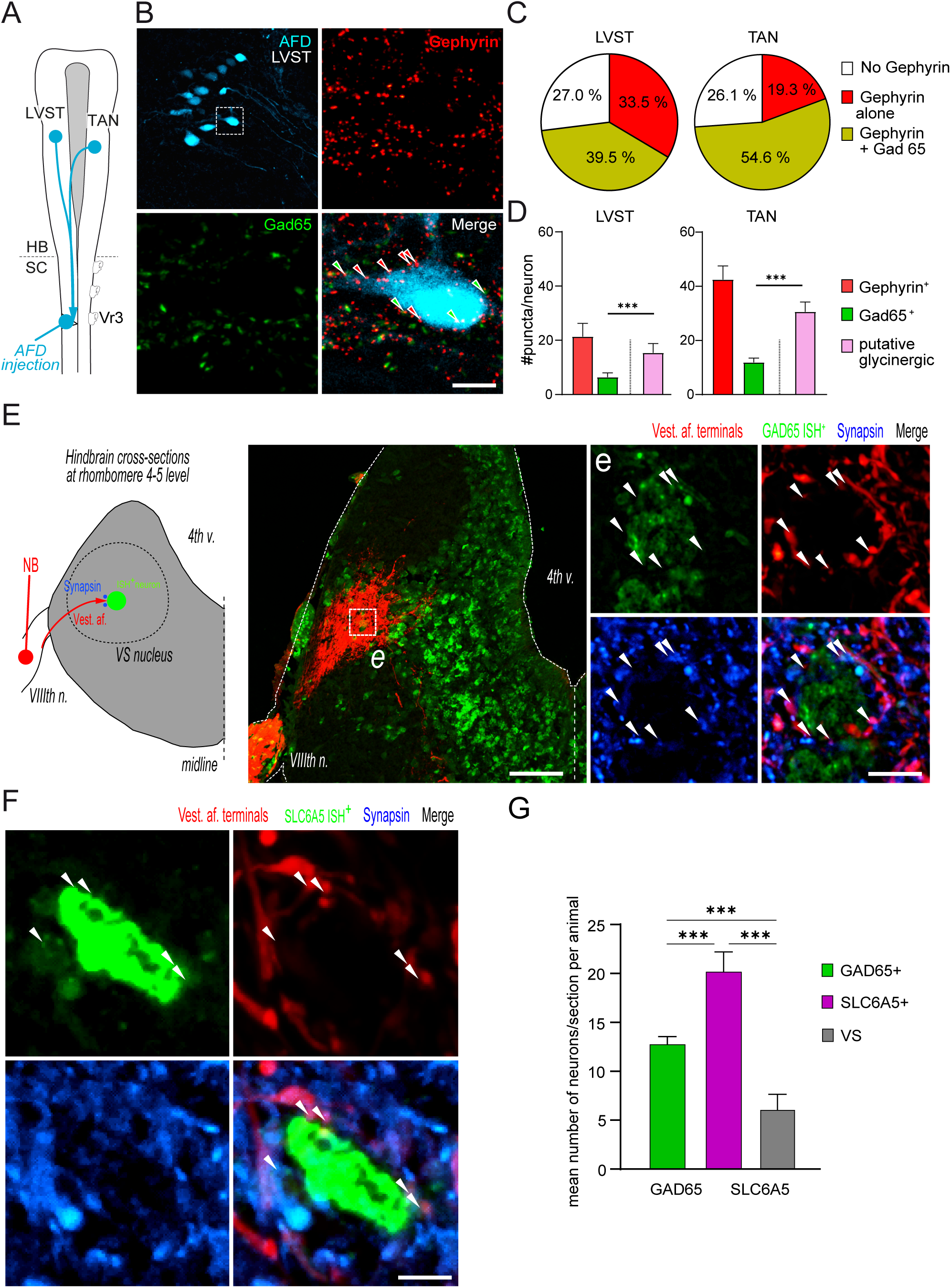
Local inhibitory network in vestibulospinal nuclei of the *Xenopus* tadpole. **A-D**: Vestibulospinal neurons display numerous inhibitory synapse markers. A: Experimental design for VS neuron retrograde labeling. B: Immunofluorescence detection of both gephyrin (red) and Gad65 (green) protein was performed in all labeled VS-containing brainstem slices. C: Distribution of identified VS populations according to the type of inhibitory synapses they display. D: Distribution of gephyrin- and Gad65-positive puncta on identified VS neurons, indicating the mean number of GABA-A and likely glycinergic input synapses. **E-G**: FISH identification of inhibitory secondary vestibular neurons in VS nuclei. E: Experimental design. F: GABAergic neurons in both VS nuclei were monosynaptically contacted by vestibular sensory afferents, as illustrated by the colocalization of GAD65 (green), neurobiotin (NB; red) and synapsin (blue) signals. F: Glycinergic neurons in both VS nuclei were monosynaptically contacted by vestibular sensory afferents, as illustrated by the colocalization of SLC65 (green), neurobiotin (NB; red)) and synapsin (blue) signals. G: Relative distribution of VS, GAD65^+^ and SLC65^+^ neurons in the VS nuclei.

Previous studies in frogs suggested that VS neurons are surrounded by inhibitory neurons (see Rössert & Straka, 2011). Using the FISH technique, we investigated the distribution of inhibitory neurons in the brainstem of *Xenopus* tadpoles, using either GAD65 or SLC6A5 probes to respectively characterize GABAergic and glycinergic cell bodies. Beforehand, neurobiotin (NB) was injected either in a VIII^th^-nerve root to label vestibular sensory afferents on one side (Fig. 1E-F), or in proximal spinal segments to label VS neurons in order to delineate anatomically the LVST and the TAN nuclei. Both GAD65 and SLC6A5-positive cell bodies were observed (n=7 animals) in comparable and abundant proportions in the two vestibulospinal nuclei, to be contacted directly by vestibular afferents (Fig. 1E-F), demonstrating they were indeed secondary vestibular neurons. Together in the two CVNs, we found very small proportions, about 3.9% (10/257) and 1.0% (18/575), of identified VS neurons expressing either GAD65 or SLC6A5 markers, respectively (not illustrated). However, because their impact would be exclusively spinal, we did not consider further these two specific populations of inhibitory neurons in the present study. Besides such inhibitory VS neurons, GAD65-positive and SLC6A5-positive cell bodies were found in comparable proportions in the two VS nuclei and represent significantly larger neuronal populations (ANOVA; p<0.001), with respectively 12.8±0.3 neurons per slice (n=26 slices, 5 tadpoles) and 20.2±0.9 neurons per slice (n=19 slices, 5 tadpoles; Fig. 1G) than VS neurons (6.2±0.7 neurons per slice). The greater proportion of glycinergic compared to GABAergic neurons (Fig. 1 G; p<0.001) mirrored the higher number of likely glycinergic input synapses onto VS neurons reported above.

We then investigated whether neurons of vestibular commissural system, which was shown to exert an overall inhibition onto contralateral CVNs in the VOR context (Malinvaud *et al*., 2010), expressed any marker of inhibition. Using GABA immunodetection we found a restrained population of retrogradely-marked VC neurons showing double labelling (12.1%; n=190 VC neurons; Fig. 2A-B), suggesting that very little number of VC neurons were inhibitory. The inhibitory nature of a small part of the VC neurons (n=326) was later confirmed in another set of animals where FISH on NB-filled VC neurons revealed that only 10.8% and 4.6% of VC cell bodies displayed GAD65 or SLC6A5 signal, respectively. We then investigated the neuronal targets of VC neurons within the contralateral VS nuclei. First, we identified Gad65-immunoreactive VC neuron terminals in the very close vicinity of retrogradely-labelled VS neuron cells bodies (Fig. 2C-D), strongly suggesting that GABAergic VC neurons made direct synapses onto contralateral VS neurons. In addition, using FISH and NB-filled VC terminals (Fig. 2E-G) we observed numerous synapsin-identified synaptic contacts between VC terminals and both GAD65^+^ (Fig. 2F) and SLC6A5^+^ cell bodies (Fig. 2G). Taken together, our observations in Xenopus tadpole supported a majorly inhibitory action of the VC system onto contralateral VS neurons, through both by direct GABAergic inhibition and indirect, local GABA and/or glycine interneuron-mediated inhibition of VS neurons.

**Figure 2:**
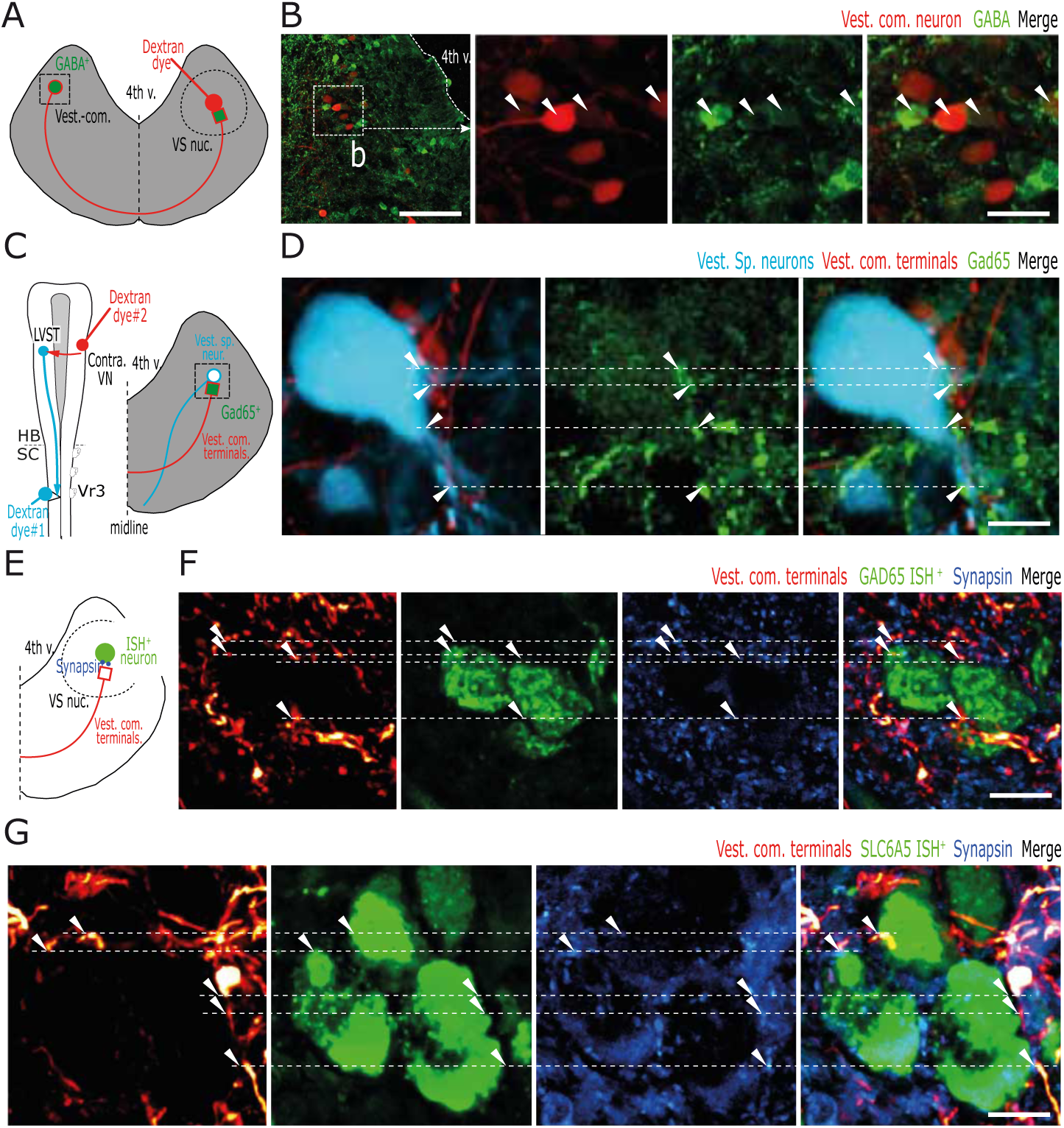
Commissural inhibition in vestibulospinal nuclei of the *Xenopus* tadpole. **A-B**: Identification of GABAergic VC neurons. A: Experimental design. B: GABA immunolabeling (green) could be observed in a small part of retrogradely marked VC neurons (red). **C-D**: GABAergic VC neurons project onto contralateral VS neurons. C: Experimental design. D: Close appositions of Gad65^+^ (green) VC terminals (red) and VS cell body and dendrites (blue). **E-G**: VC neurons project onto inhibitory neurons in contralateral VS nuclei. E: Experimental design. F: Colocalization of GAD65 FISH (green), VC terminals (red) and synapsin (blue), indicating direct synapses between VC neurons and contralateral GABAergic secondary vestibular neurons. G: Colocalization of SLC65 FISH (green), VC terminals (red) and synapsin (blue), indicating direct synapses between VC neurons and contralateral glycinergic secondary vestibular neurons.

### GABAergic and glycinergic postsynaptic potentials in identified VS neurons

Taken together, the above results strongly suggested that VS neurons received massive inhibitory inputs from brainstem GABAergic and glycinergic interneurons, either locally or contralaterally (VC interneurons) distributed. We performed whole-cell patch-clamp recordings in both LVST and TAN nuclei from identified VS neurons, priorly labeled with retrograde RDA injected in the rostral spinal segments. We confirmed that VS neurons received massive inhibitory inputs from brainstem GABAergic and glycinergic interneurons (Fig. 3). In current clamp configuration, we characterized the occurrence, frequency, amplitude, duration and pharmacology of spontaneous IPSPs in the permanent presence of glutamate-receptor antagonists CNQX (10 µM) and APV (20 µM).

**Figure 3:**
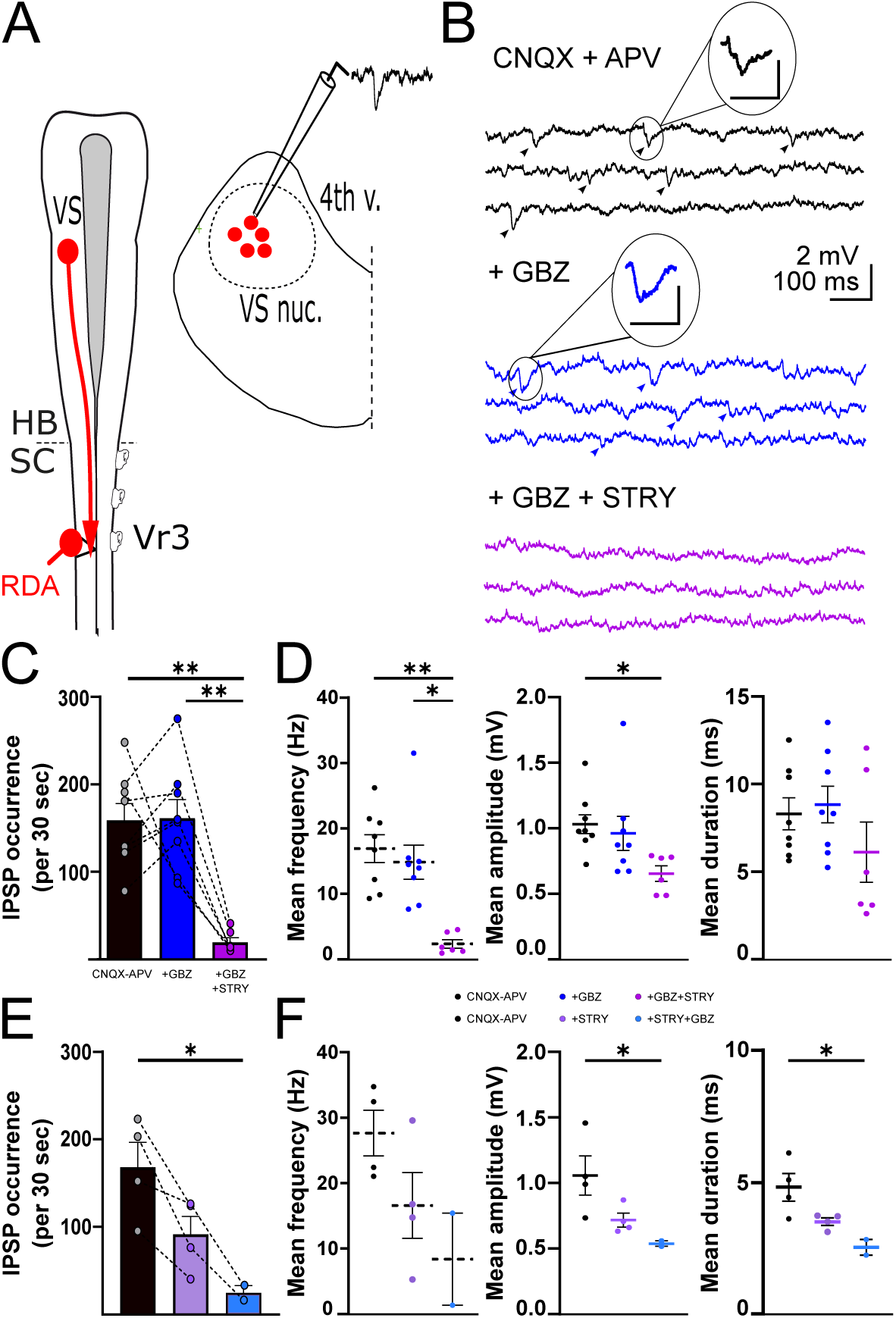
Patch-clamp characterization of inhibitory postsynaptic potentials in vestibulospinal neurons of the *Xenopus* tadpole. **A**: Experimental design. **B**: Spontaneous IPSPs (arrowheads) were recorded from identified VS neurons in control condition (black) and after gabazine (+GBZ, blue) and subsequent strychnine (+GBZ+STRY, purple) addition to the perfusion bath. GABA immunolabeling (green) could be observed in a small part of retrogradely marked VC neurons (red). Three consecutive episodes are illustrated for each condition. Insets illustrate IPSP doublets (scalebars: 1 mV, 10 ms). **C-D**: Occurrence (C) and parameters (D) of IPSPs in control (black) and after gabazine (blue) and subsequent strychnine (purple) perfusion. **E-F**: Same arrangement as C-D but with strychnine perfused first (light purple) and gabazine added subsequently (light blue).

In these conditions, IPSPs occurred spontaneously at 17.0±2.1 Hz (Fig. 3B black traces, and black symbols in figure graphs; n=12) with a small amplitude (1.03±0.07 mV) and long duration (8.31±0.90 ms). In a first set of experiments, the GABA-A receptor antagonist gabazine (1 µM) was added first to the perfusion bath. Surprisingly, this did not affect significantly the occurrence of spontaneous IPSPs which remained abundant, with a mean frequency (14.9±2.6 Hz), amplitude (0.96±0.13 mV) and duration (8.84±1.04 ms) similar as controls (Fig. 3B blue traces, and dark blue symbols Fig. 3C-D; n=8). In contrast, adding the glycinergic receptor antagonist strychnine (0.1 µM) to the CNQX-APV-gabazine medium (Fig. 3B purple traces, and purple symbols in Fig. 3C-D; n=6) dramatically reduced the occurrence of spontaneous IPSPs (p<0.01), the frequency of which dropped to 2.4±0.6 Hz (p<0.01), as well as IPSP mean amplitude (0.65±0.06 mV; p<0.05). These results suggested that VS neurons received mostly, if not only spontaneous glycinergic IPSPs since the lack of gabazine effect on IPSP occurrence and physiological parameters supposed that there were no direct GABA-mediated inputs on VS neurons. Because it was on striking contradiction with our anatomical results, we hypothesized that gabazine perfusion could have unmasked additional glycinergic IPSPs. Therefore, in a second set of experiments (Fig. 3E-F; n=4) we reversed the order of inhibitory antagonist application: strychnine applied first strongly decreased the occurrence of spontaneous IPSPs (from 168±29 in control to 92±21 under strychnine), with a significant tendency to reduce the mean frequency, amplitude and duration (p=0.057; Fig. 3E-F, light purple symbols). After adding gabazine to the strychnine perfusion almost removed all IPSPs, the remaining exhibiting a significant reduction in both amplitude and duration (Fig. 3E-F, light blue symbols).

### GABAergic and glycinergic modulation of GVS-evoked reflexes

The above neuroanatomical and functional data strongly supported the hypothesis that VS neurons in both LVST and TAN nuclei are modulated by GABA and glycine inhibitory inputs. Then, we investigated how the perfusion of inhibitory antagonists onto the brainstem could affect the integration of vestibular sensory inputs and the resulting spinal reflexes.

Spinal ventral root activity was recorded in response to horizontal canal activation with GVS in semi-intact preparations, while either gabazine (1 µM; n=10) or strychnine (0.1 µM; n=10) was restrictively perfused on the brainstem (Fig. 4). As previously reported (Barrios *et al*., 2024), GVS evoked various forms of spinal responses along the spinal cord that were all characterized by left-right alternation phase-locked on the GVS cycle. Because the most stable responses were recorded from the initial tail segments where postural responses corresponded to the maximal tail curvature (Bacqué-Cazenave *et al*., 2022; Barrios *et al*., 2024), activities from left and right ventral roots #11 were used to illustrate the spinal cord behavior: maximal motor response of left-side spinal roots occurred at the peak of right-side horizontal canal stimulation, and conversely (Fig. 4A and D).

**Figure 4:**
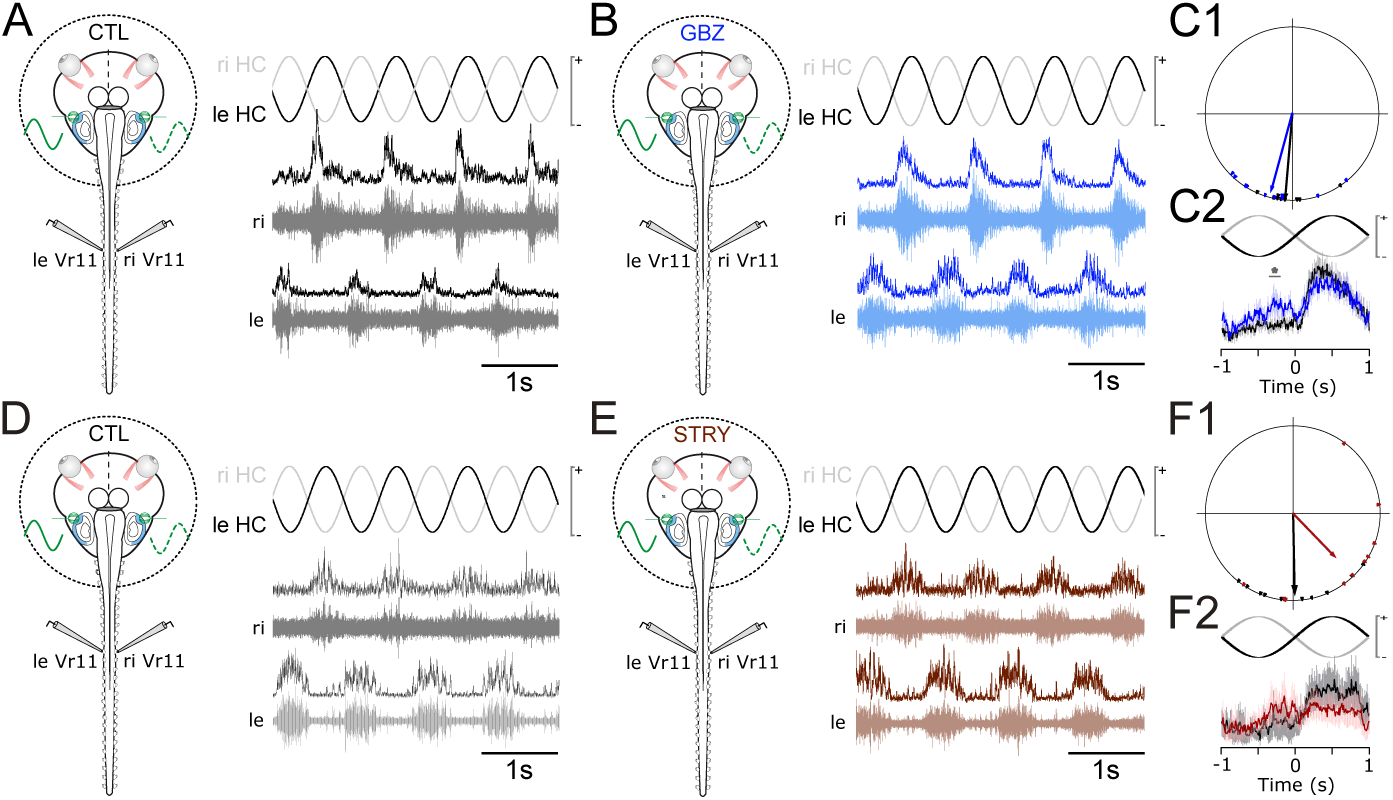
Effects of brainstem-restricted perfusion of inhibitory antagonists on the timing of GVS-evoked spinal reflexes in the *Xenopus* tadpole. **A-C**: Gabazine (GBZ) effects. A: Control spinal reflex response evoked by sinewave stimulation of horizontal canal cupula on both sides. B: Reflex evoked under gabazine perfusion. C: Circular distribution (C1) of spinal burst peaks and mean discharge rate (C2; arbitrary units) in control (black) and under gabazine (blue). Note the appearance of a secondary little peak during the non-preferred phase of the GVS (* p<0.05). **D-F**: Strychnine (STRY) effects. Same arrangement as A-C. Note the loss of coherence in the phase distribution (F1) and the spreading of the ‘peak’ response (F2) under strychnine.

Apparently, neither gabazine (Fig. 4B) nor strychnine (Fig. 4E) perfusion on brainstem dramatically altered the overall GVS-evoked spinal reflex, which remained characterized by left-right alternation locked on the GVS phase (compare Fig.4A and 4B, and 4E and 4F, respectively). However, although mean phase of the response was not significantly modified by gabazine (Watson-Williams F-test; p=0.294; Fig. 4C1), the temporal structure of the response changed, as illustrated by the small magnitude but significant increase in mean discharge rate during the unpreferred GVS phase (i.e., the phase which usually does not evoke reflex activity; Fig. 4C2). Conversely, strychnine caused a large advance in the spinal discharge timing relative to the GVS cycle (Watson-Williams F-Test; p<0.05; Fig. 4F1), implying that glycine-receptor blockade caused the brainstem sensory-motor network to be more excitable. In addition, the timing coherence of the reflex responses was reduced since the mean vector length dropped from 0.87 in control to 0.69 under strychnine (respectively, black and red vectors in Fig. 4F1), and the response showed a much wider time course and started before the preferred GVS phase (Fig. 4F2). Taken together, there results demonstrate that blocking vestibular-related inhibitory systems affects the temporal accuracy of the GVS-evoked spinal responses. Conversely, brainstem GABA-A and glycine inhibitions are responsible for the precise timing of the vestibular sensorimotor processing.

We then considered the time course of repetitive spinal bursts throughout GVS repeated cycles under gabazine (Fig. 5A-C) and strychnine (Fig. 5D-F) perfusion on the brainstem. Whereas strychnine did not significantly affect spinal reflex repetition over GVS cycles, gabazine significantly changed the strength of spinal responses, measured as the area under the response curve (Fig. 5B; p>0.001). Interestingly, brainstem bath-perfusion with gabazine had antagonistic effects on the first reflex burst compared to the following ones: it reliably enhanced the response to the first GVS cycle (Fig. 5C1) and reduced significantly the spinal responses from the seventh to the last (Fig. 5C2) GVS cycle. These later results suggested distinct modes of action between GABA and glycine inhibitory systems, and further suggested a more complex role of GABA-A-mediated inhibition in the modulation of vestibulo-spinal reflexes in *Xenopus* tadpoles.

**Figure 5:**
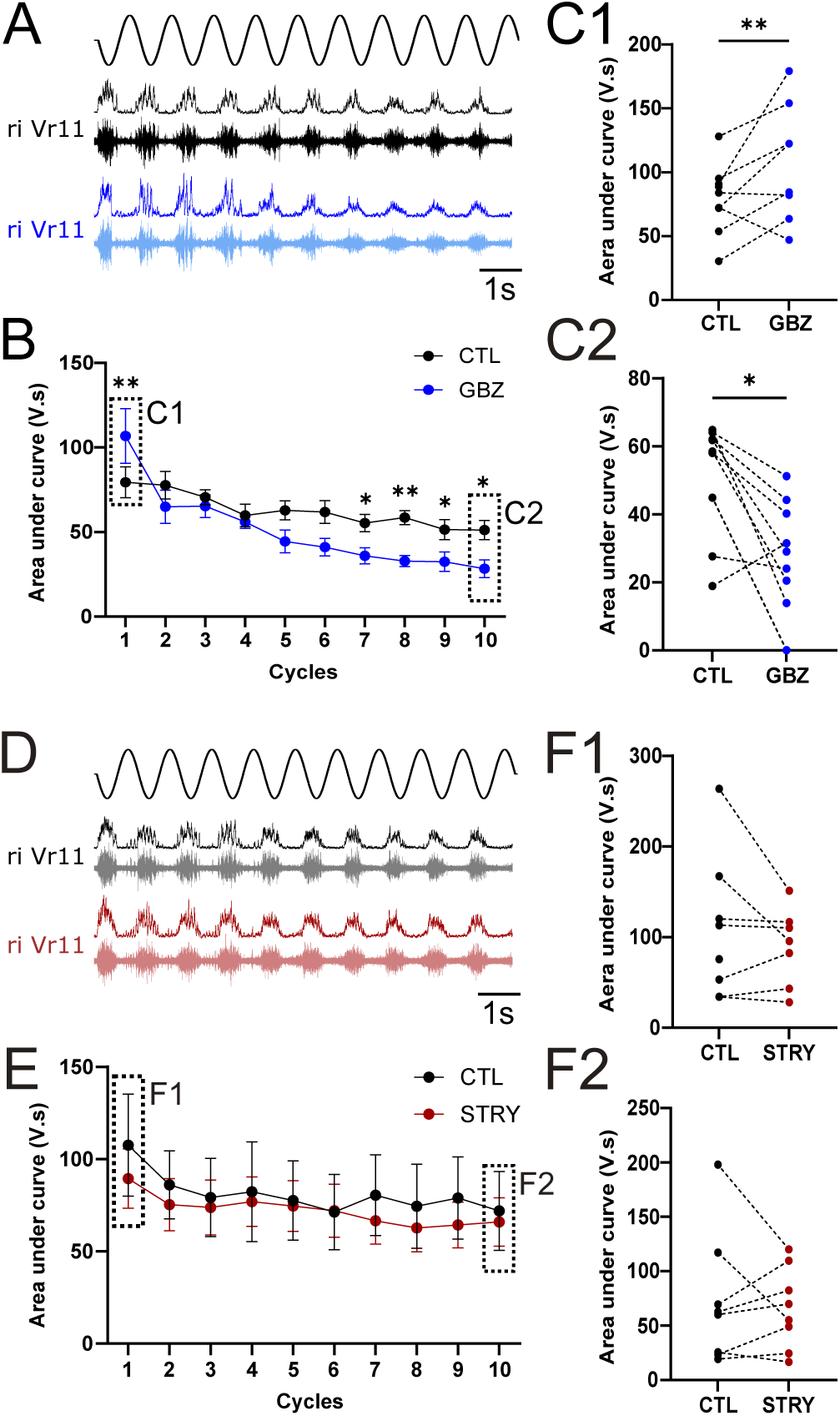
Effects of brainstem-restricted perfusion of inhibitory antagonists on the magnitude of GVS-evoked spinal reflexes in the *Xenopus* tadpole. **A-C**: Gabazine (GBZ) effects. A: GVS-evoked spinal response throughout the 10-cycle repeated stimulation in control (black and under gabazine perfusion (blue). Both the raw traces (bottom) and the integrated signals (top) are illustrated. B: Evolution of the response magnitude (measured as the mean area under the integrated curve). C: First (C1) and last (C2) mean areas in control (black) and under gabazine (blue). Note the distinct effects between the first and later GVS cycles (** p>0.01; * p<0.05). **D-F**: Strychnine (STRY) effects. Same arrangement as A-C.

### GABAergic and glycinergic modulation of VS neuron stimulation-evoked reflexes

Previous works in the terrestrial frog showed that semicircular canal stimulation evoked both excitation and inhibition in CVN unidentified neurons (Straka & Dieringer, 2000; Biesdorf *et al*., 2008). It was thus most likely that perfusion of inhibitory receptor antagonist in our GVS experiments also affected this well described component of the sensory-motor integration. Because our goal was to focus on inhibition on VS neurons, spinal motor responses were thereafter evoked by direct electrical stimulation of either LVST or TAN nucleus while the brainstem was bathed in control, gabazine (1 µM) or strychnine (0.1 µM) perfusion.

As recently described (Barrios *et al*., 2024) single-shock electrical stimulation of either LVST (Fig. 6A-D, 6I-M) or TAN (Fig. 6E-H, 6N-R) in control conditions triggers both fast and slow compound bursts recorded from spinal ventral roots. Because long delay responses depended on spinal network rather than on VS excitability, we focused our analysis on the fast responses that occurred between 3 and 20 ms in control conditions (Fig. 6; LVST: panels B1 and J1; TAN: F1 and O1), which engaged monosynaptic rather than oligosynaptic connections, and thus reflected better how much VS neurons were activated by the delivered electrical pulses. In these conditions, gabazine perfusion on the brainstem did not affect the response delay (not illustrated) but dramatically reduced the ability of the stimulus to trigger fast spinal responses both for LVST (Fig. 6B2-C; n=10; Wilcoxon test, p<0.05) and TAN stimulation (Fig. 6F2-G; n=10; p<0.01). In addition, the strength of the remaining reflex responses (measured as the area under the response curve) was also reduced for both LVST (Fig. 6D; n=10; CTL: 3.26±0.55 mV.s; GBZ: 1.42±0.45 mV.s; Wilcoxon test, p<0.05) and TAN stimulation (Fig. 6H; n=10; CTL: 3.49±0.80 mV.s; GBZ: 0.81±0.31 mV.s; p<0.01). This unexpected result suggested that blocking GABA-A-mediated inhibition might allow another local inhibitory system to express freely, likely the glycinergic one.

**Figure 6:**
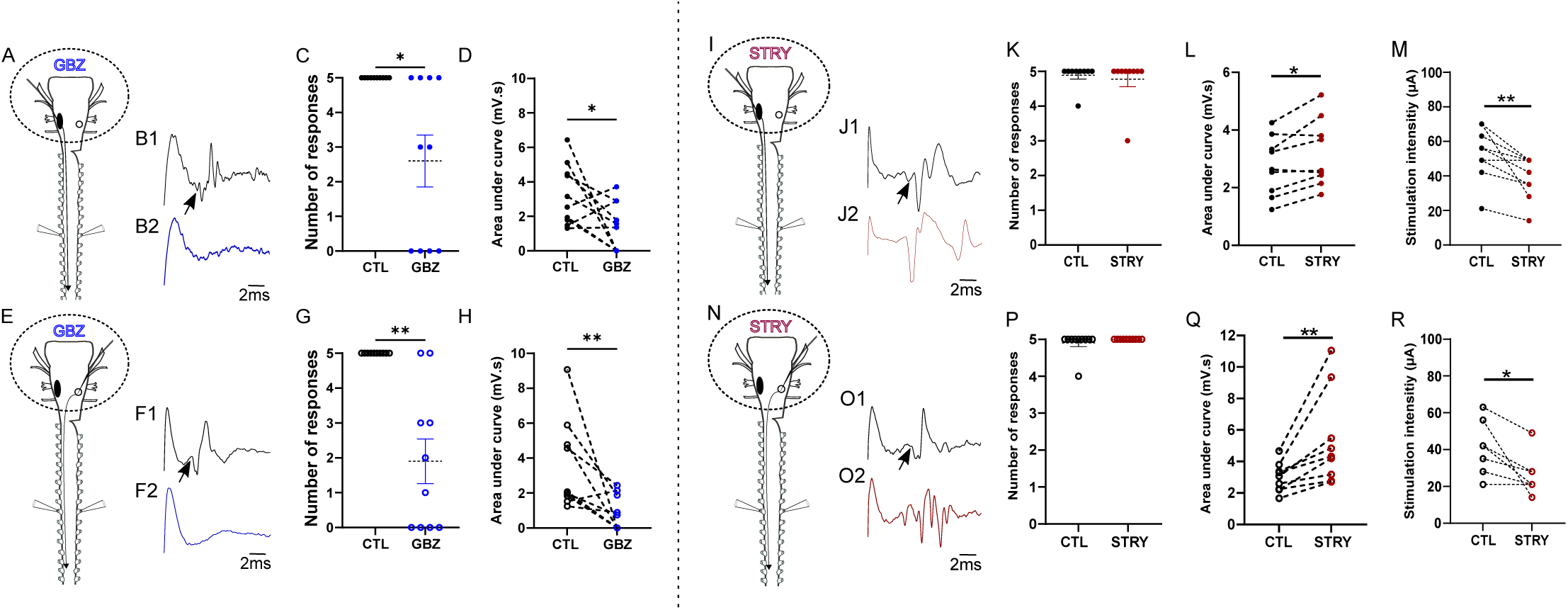
Effects of brainstem-restricted perfusion of inhibitory antagonists on CVN electrical stimulation-evoked reflexes in the *Xenopus* tadpole. **A-H**: Gabazine (GBZ) effects on spinal responses evoked by LVST (A-D) or TAN (E-H) direct stimulation. A: Experimental design for LVST. B: Samples of spinal fast responses in control (black) and under gabazine (blue). Arrowhead point to the response onset. C: Mean occurrence of spinal fast responses over 5 repeated stimulations (* p<0.05). D: Mean areas under the integrated curve of the response (* p<0.05). E-H: Same arrangement as A-D, for TAN stimulation. **I-R**: Strychnine (STRY) effects. Same arrangement as A-H.

Contrasting with gabazine, strychnine perfusion on the brainstem had no effects on the occurrence of spinal responses at the control threshold stimulation of LVST (Fig. 6J2-K; n=9; Wilcoxon test, p>0.99) or TAN nucleus (Fig. 6O2-P; n=9; p>0.99). Though, the strength of the evoked spinal discharge increased significantly in response to both LVST (Fig. 6L; n=9; CTL: 2.74±0.34 mV.s; STRY: 3.18±0.39 mV.s; Wilcoxon test, p<0.05) and TAN stimulation (Fig. 6Q; n=9; CTL: 2.89±0.31 mV.s; STRY: 5.32±0.98 mV.s; p<0.01). In addition, the threshold intensity required to evoke a spinal response was always lower than in control saline for LVST (Fig. 6M; n=9; control: 52.9±5.1 µA; strychnine: 38.9±4.1 µA; Wilcoxon test, p<0.01) as well as for TAN stimulation (Fig. 6R; n=9; control: 38.9±4.8 µA; strychnine: 25.7±3.3 µA; p<0.05), which demonstrated a strong increase in VS neuron excitability after blocking brainstem glycinergic inhibition.

Taken together, these stimulation experiments suggested that VS neuron activity was tonically depressed by inhibitory glycinergic inputs, whereas GABA-A neurotransmission rather regulated the expression of such a tonic glycinergic inhibition.

### Role of vestibular commissural neurons in vestibular-induced spinal reflexes

In terrestrial frogs, VC inhibition is mediated by both GABAergic and glutamatergic secondary vestibular neurons projecting into contralateral CVNs (Malinvaud *et al*., 2010), and its implication in the vestibulo-ocular reflex has been well documented (Straka & Dieringer, 2004). Because brainstem inhibitory networks play major roles in organizing the vestibular-induced spinal reflexes (see above), we thus investigated whether brainstem commissural connections might regulate GVS-evoked spinal reflexes as well (Fig. 7). To this end, the rostral population of VC connections was severed by splitting the brainstem along the midline between rhombomeres 1 and 3 (n=11), in order to keep intact all the crossing vestibulospinal axons originating in the TAN nucleus (Straka *et al*., 2001). In most preparations (8/11), the classical GVS-locked, left-right alternating spinal pattern recorded in control conditions (Fig. 7A-B) got turned into a bilaterally synchronous reflex occurring at either one phase (Fig. 7C-D) or the two phases (not illustrated) of the GVS cycle after commissural disruption. Such a dramatic modification of the response pattern demonstrated that VC integrity is required for accurate postural reflexes to be produce.

**Figure 7:**
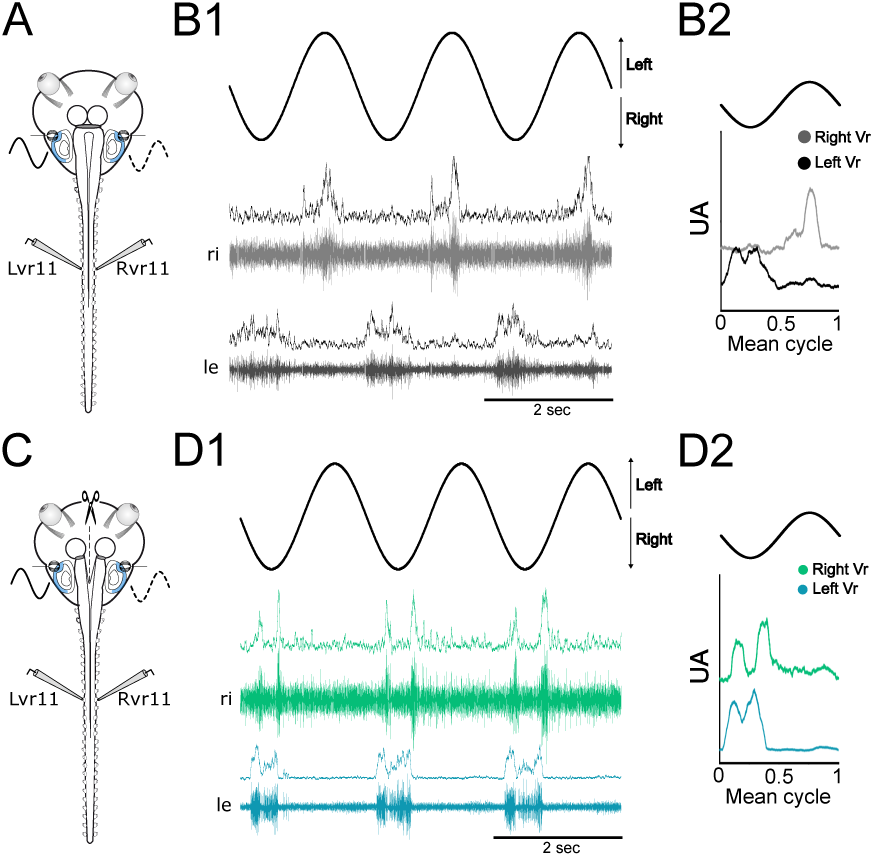
Effects of rostral vestibulo-commissural pathway disruption on GVS-evoked spinal reflexes in the *Xenopus* tadpole. **A-B**: Control response. A: Experimental design. B: Sample left and right-side GVS-evoked activity recorded from the eleventh ventral root (B1), and corresponding mean discharge rate (B2; arbitrary units). **C-D**: After rostral VC axon transection. Same arrangement as A-H.

In another set of experiments, either the LVST (n=13; Fig. 8A-D) or TAN nucleus (n=12; Fig. 8E-H) was directly stimulated before and after severing the rostral commissural pathways, to get rid of the sensory influence and analyze specifically the VC control onto VS neurons. Disrupting the VC connections could have three distinct consequences, whatever the stimulated VS nucleus: either no significant effects on the evoked spinal response (not illustrated), or a reduction (Fig. 8B and F, top middle traces) or complete loss of the fast spinal response (Fig. 8B and F, bottom middle traces). Indeed, in more than half of the preparations the ability of a LVST (Fig. 8C; ANOVA, p<0.001) or TAN (Fig. 8G; ANOVA, p<0.01) threshold stimulation to evoke fast spinal responses was strongly reduced. Interestingly, adding gabazine to the perfusion bath after VC cut did not affect more the ability of LVST stimulation to evoke spinal responses (Tukey’s post-ANOVA test; p>0.05), while it potentiates the disruptive effect of VC pathway cutting on TAN stimulation efficiency to trigger spinal responses (Tukey’s post-ANOVA test; p<0.05). Then, the strength of spinal responses was analyzed only for the stimulation sites that exhibited reduced efficacy after commissural interruption (Fig. 8D, LVST: 9/13; Fig. 8H, TAN: 6/12), and a significant decrease was observed for both CVN (green bars) compared to control (empty bars). Interestingly, adding gabazine (1 µM) after VC cut did not affect further the evoked response (dark green bars). Taken together, these results suggested that the GABA-A control over VS excitability might occur, at least for a large part, through the activation of the VC connections.

**Figure 8:**
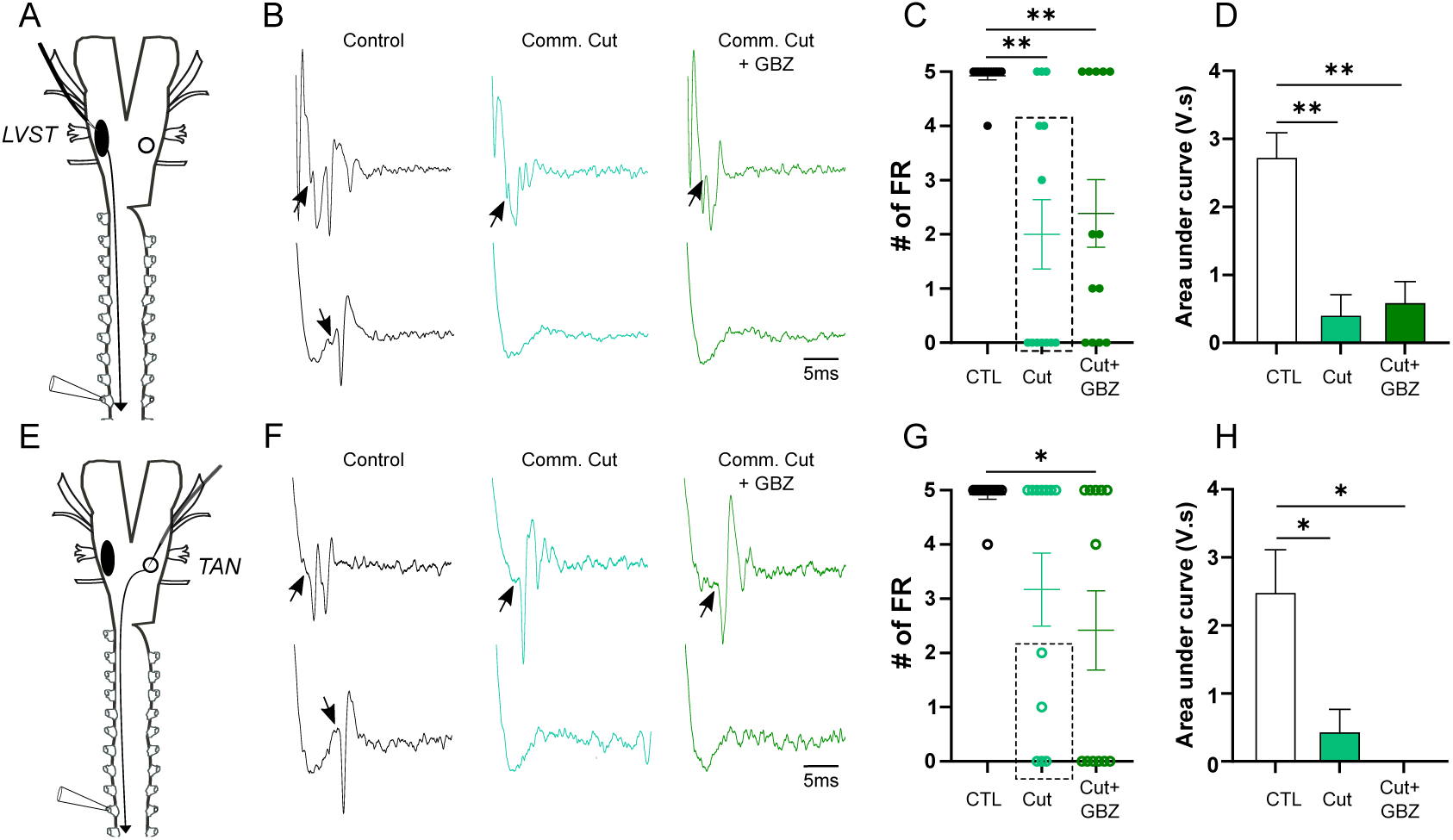
Effects of rostral vestibulo-commissural pathway disruption on CVN electrical stimulation-evoked reflexes in the *Xenopus* tadpole. **A-D**: LVST-evoked responses. A: Experimental design. B: Sample spinal responses to cut-sensitive (bottom) and insensitive (top) stimulation sites in the LVST, in control (black), and after commissural cut (light green) and subsequent gabazine perfusion (dark green). Arrowheads point to response onsets. C: Mean response occurrence over 5 consecutive stimulations. D: Mean response magnitude (measured as the mean area under the integrated response curve; * p<0.05) in preparations affected by the commissural cut (doted box in C). **E-H**: TAN-evoked responses. Same arrangement as A-D.

## Discussion

VS reflexes are essential in vertebrates to ensure accurate body and head orientation in space at rest and during locomotion. Whereas the excitatory pathways had been well described in a large variety of animal models, little attention was paid to the surrounding inhibitory networks and their functional role in postural reflexes. Here, focusing on the vestibulospinal control of postural reflexes we provide a series of evidences that, taken together, strongly argue in favor of a pivotal role of vestibular-related brainstem inhibitory networks in orchestrating vestibular-induced spinal responses. First, numerous vestibular-related local GABAergic and glycinergic neurons exist within CVNs, and inhibitory inputs, partly commissural, impact onto VS neurons. Second, the blockade of one or the other inhibitory component modified the timing and/or accuracy of the GVS-evoked spinal reflex. Third, interruption of the rostral VC connections, which net effect is known to be inhibitory on contralateral CVNs in frogs (Straka & Dieringer, 2004; Malinvaud *et al*., 2010), had dramatic consequences on the reflex pattern. Finally, we demonstrated a complex interaction between GABA-A and glycine-mediated inhibition onto VS neurons, which regulates VS neuron ability to trigger spinal reflexes, and likely involves the VC pathways.

The presence of inhibitory neurons in CVNs was demonstrated long ago in various vertebrates (*e.g.*, Holstein *et al*., 1996). They are activated directly or indirectly by vestibular afferents, and act either locally (ipsilateral to the vestibular input) or on the contralateral CVNs through commissural connections. This organization was essentially described for the VOR loop in both mammals (reviewed in Gliddon *et al*., 2005) and amphibians (reviewed in Straka & Dieringer, 2004), but remained so far unknown in vestibulospinal circuits. This study shows that a comparable organization of brainstem circuits also exists in VS pathways, with the implication of both local and commissural vestibular inhibitory neurons. In frogs, vestibular afferents were shown to trigger both monosynaptic excitatory and disynaptic inhibitory PSPs in ipsilateral unidentified CVN neurons and disynaptic IPSPs in contralateral ones (Straka & Dieringer, 2004). The combination of such inputs was proposed to regulate the timing of single CVN neuron discharge in response to a vestibular sensory input. Along that line, we investigated the functional consequences of such inhibitory neurons in VS reflexes. We demonstrated here that both GABA and glycine-mediated inhibitions tuned the phase of the spinal response to GVS applied on horizontal canals (Fig. 4). The timing of the spinal reflex likely was the consequence of a neuronal processing within CVNs, integrating horizontal canal sensory signals in a given context of excitation/inhibition balance (as previously reported in unidentified CVN neurons). We provided additional evidences that the two inhibitory systems acted distinctly on VS neuron excitability. A GABA-A antagonist reduced the strength of the spinal reflex while glycine-receptor blockade favored the reflex expression (Fig. 6). As a result, whereas GABA-A receptor blockade slightly changed the temporal structure of the spinal reflex response, glycinergic blockade in contrast advanced the response phase (Fig. 4). This difference might be explained by the interaction that seemed to exist between the two neurotransmitter systems. Indeed, our patch-clamp recordings clearly demonstrated that VS neurons spontaneously exhibited sustained IPSPs of both GABA-A and glycine origins (Fig. 3). In addition, whereas gabazine primarily applied did not affect significantly the number of spontaneous IPSPs, strychnine applied first did not either suppress all IPSPs, and the remaining IPSPs almost disappeared after gabazine addition (Fig. 3E). This can be interpreted as GABA acting both on VS neurons and presynaptic glycinergic CVN interneurons. With such a network organization, gabazine would block GABA-A IPSP in VS neurons but also release additional glycinergic IPSPs in VS neurons. Conversely, strychnine would only prevent glycinergic IPSPs while GABA-A ones would remain until additional application of gabazine. Actually, GABA-A-mediated control over glycinergic neuron activity had already been described in rat spinal cord, where presynaptic GABA-A receptors on glycinergic neuron terminals altered glycine release onto dorsal neurons (Jang *et al*., 2002). Such a presynaptic control may also exist in CVNs of *Xenopus* tadpoles, although we cannot exclude a somatic GABA-A-mediated inhibition of glycinergic neurons as well.

Besides the local inhibitory neurons, inhibition on CVN neurons may also arise from VC neurons and the cerebellum (see Gliddon *et al*., 2005). Although the cerebellum is reduced to a single lamella crossing the midline above the rhombencephalon-mesencephalon border in amphibians, it sends GABA-A signals towards CVN neurons, and their suppression was shown to increase the VOR gain in adult terrestrial frogs (Straka & Dieringer, 2004) and prevent VOR homeostatic plasticity in *Xenopus* tadpoles (Dietrich & Straka, 2016). In our experiments GABA-A receptor antagonist impaired both the ability of CVN stimulation to generate a spinal response and the magnitude of the evoked motor burst (Fig. 6A-H). This suggests that GABA neurotransmission tonically favors VS neuron excitability, likely through the inhibition of presynaptic glycinergic neurons (see above). Such a mechanism thus contrasts with the previously described cerebellar modulation of VOR and suggests that, in *Xenopus* tadpoles, the cerebellum is not involved in the gain of the horizontal canal-induced reflex. Another possibility would be that the cerebellar modulation targets specifically vestibular sensory inputs. Yet, we found that gabazine perfused on the brainstem during GVS induced a little phase retardation of the reflex (Fig. 4C1), demonstrating that conversely, the GABAergic control favored VS sensory-triggered motor responses.

VC neurons bridge lateralized vestibular sensory inputs to contralateral CVNs. Although using both excitatory and inhibitory neurotransmissions the net impact of such crossing neurons is inhibitory on their contralateral target nuclei (Barmack, 2003; Straka & Dieringer, 2004). In grass frogs, for instance, the VC system comprises both glutamatergic and GABAergic neurons, the activation of which evokes both large amplitude EPSPs and small amplitude IPSPs in contralateral CVN neurons, mostly with a monosynaptic-compatible latency (Malinvaud *et al*., 2010). In the *Xenopus* tadpole (as was shown also in the perinatal mouse; Dubois *et al*., 2022), we found only a small proportion of commissural neurons expressing GABA markers (half the number reported in the adult grass frog) and another small population exhibiting glycinergic markers, suggesting that, as in other vertebrates (Gliddon *et al*., 2005) the large majority of VC neurons is excitatory. This suggested further that the majority of such excitatory VC neurons project onto local inhibitory neurons within the contralateral CVNs to achieve commissural inhibition. Such organization was recently reported in mice where GABAergic neurons in the medial vestibular nucleus were found to receive direct inputs from contralateral VC neurons (Kong *et al*., 2024). We found that severing the rostral VC pathway could have similar effects as gabazine perfusion on CVN stimulation-evoked spinal responses (Fig. 8), suggesting that either excitatory VC neurons in the *Xenopus* tadpole also project onto contralateral GABA neurons, or GABAergic VC neurons project onto contralateral glycinergic neurons, or both, to regulate VS excitability. We indeed observed direct connections between VC and local GABA and glycine CVN interneurons in the *Xenopus* tadpole (Fig. 2).

GVS experiments showed that affecting the integrity of the VC pathway had more dramatic effects on the temporal structure of VS reflexes than could have the antagonization of GABA-A or glycine-mediated inhibition (Fig. 7). The fact that only a part (the rostral one) of VC projections were severed in our experiments likely had introduced a disequilibrium between rostral and more caudal commissural influences, thus resulting in a complete loss of CVN bilateral coordination. However, note that GABA-A blockade tended to reveal an additional peak of spinal discharge during the non-preferred GVS phase (as illustrated in Fig. 4C2). In addition, such a split in the region equivalent to the pons in higher vertebrates could have also affected the global spinal homeostasis by damaging many regulating descending influences (Takakusaki *et al*., 2016; Takakusaki, 2017), including vestibular-targeted reticulospinal cells (Straka *et al*., 2001), resulting in a complete disorganization of spinal networks and their inability to produced normal reflex patterns in response to VS commands.

In conclusion, our present results demonstrate a fundamental role of brainstem inhibitory circuits in the generation of vestibular-evoked spinal reflexes in *Xenopus* tadpoles (in striking contrast with what was proposed in mice; Montardy *et al*., 2021). Together with the literature, they further point to a complex network organization (Fig. 9), involving both local and commissural neurons interacting together to orchestrate vestibular-induced postural adjustments. In summary, unilateral vestibular sensory signals activate the ipsilateral TAN VS neurons both through direct excitation and oligosynaptic disinhibition (activation of GABAergic interneurons inhibiting glycinergic interneurons) while LVST VS neurons are mostly inhibited (Fig. 9A). In parallel, vestibular afferents trigger ipsilateral VC neurons to inhibit both directly (inhibitory VC neurons) and indirectly (through activation of contralateral GABA and glycine interneurons) the TAN VS neurons on the contralateral side while activating directly (excitation) and indirectly (disinhibition) the contralateral LVST VS neurons (Fig.9B). The resultant of such excitation/inhibition balance of the various CVNs will be a stronger activation of the postural networks in the hemicord contralateral to the vestibular sensory signal and the generation of a suitable, corrective postural reflex. Notably, this organization goes farther than the simple push-pull organization proposed for the VOR since severing VC connection totally disrupt the reflex temporal structure, which was never described in the case of the VOR (e.g., Tham *et al*., 1989). In this model, the two major inhibitory neurotransmitters, glycine and GABA, are distinctly involved, the former acting rather as a rheostat controlling VS neuron excitability while the latter operate the structure of the reflex motor command.

**Figure 9:**
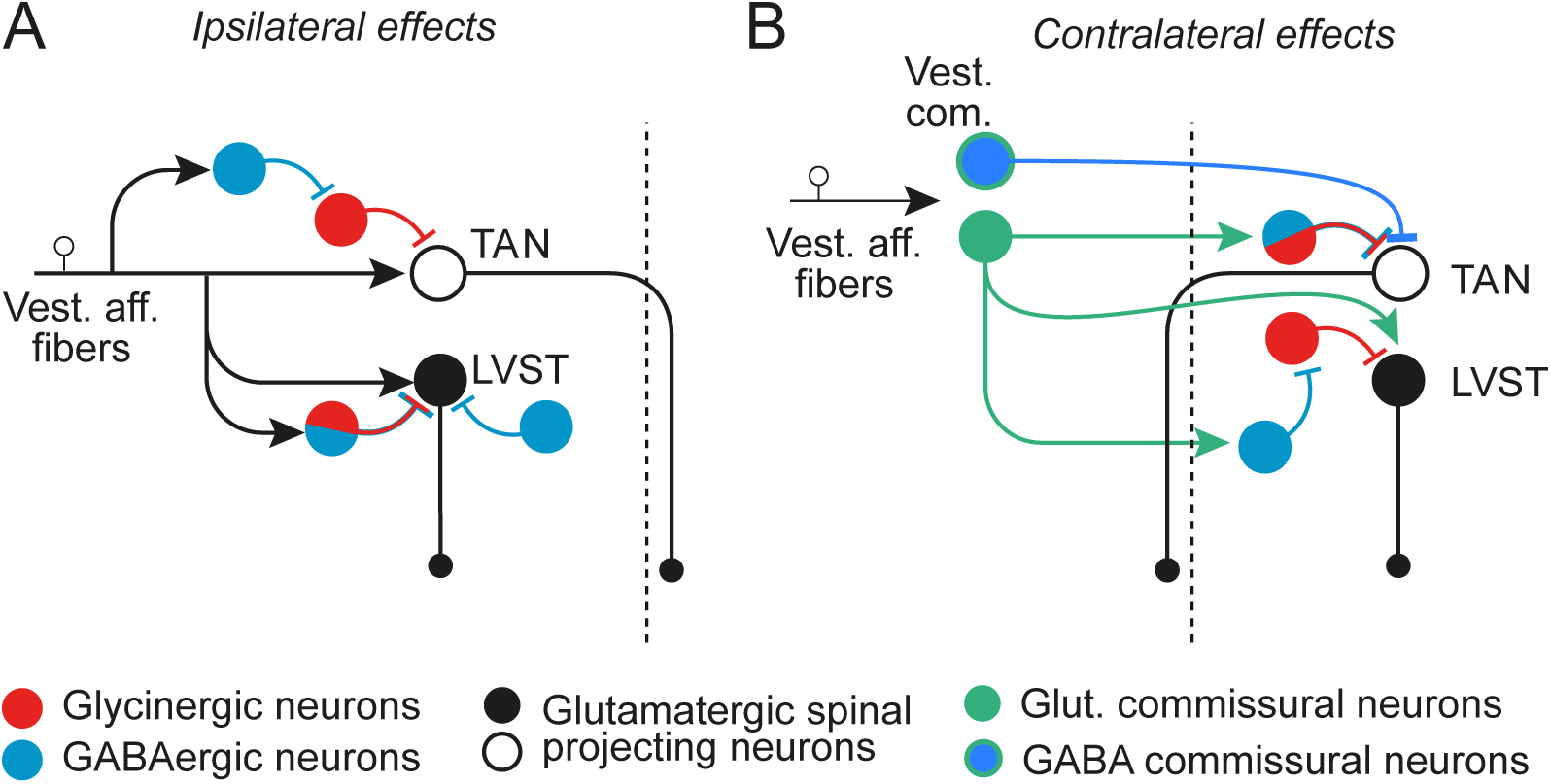
Functional organization of local and commissural inhibitory systems in vestibulospinal nuclei in the *Xenopus* tadpole. **A**: On the ipsilateral side, the distribution of afferent terminals on VS neurons and local inhibitory interneurons in CVNs results in the greater stimulation of ipsilateral TAN VS neurons, generating a stronger activation of contralateral spinal postural networks. **B**: On the contralateral side, the distribution of excitatory and inhibitory commissural terminals (arising from the sensory-activated side) onto VS neurons and CVNs inhibitory interneurons results in stronger activation of LVST VS neurons and ensuing reinforcement of the spinal networks contralateral to the vestibular sensory stimulation.

## Author contributions

All experiments were performed at the INCIA. All persons who participated in this work are listed as authors, and all authors contributed significantly to this work. DLR conceived the study. FML, PF and DLR were responsible for the operational conception of the work. LL, MP, GB, LC, MB, AD, FML and DLR were responsible for data acquisition and/or analysis. HT conceived the FISH probes. LL, FML, PF and DLR were responsible for drafting and critically revising the manuscript. All authors approved the final manuscript and agreed to be accountable for the accuracy and integrity of the work.

## Acknowledgements

This work received funding from the French government in the framework of the University of Bordeaux’s IdEx “Investments for the Future” program GPR_BRAIN_2030 (D. Le Ray & P. Fossat) and the Centre National des Etudes Spatiales (CNES – APR 2024, D. Le Ray), and was additionally supported by the Centre National de la Recherche Scientifique (CNRS).

Authors are thankful to IMN’s and INCIA’s zootechny teams for nursing the pets, to Rabia Bouali-Benazzouz (IMN) and Marie-Laure Rousseau (INCIA) for their valuable administrative help, and to Gladys Alfama and Anne-Laure Gaillard (PhyMA) for teaching LL and LC the FISH technique.

## Notes

### Competing Interest Statement

The authors have declared no competing interest.

### Summary of Updates

This revised version contains: - Additional electrophysiology experiments (CVN stimulation) - Additional neuroanatomy counting - Updated analysis and statistics - Corresponding rewriting in the results section

